# Bacterial symbionts in animal development: arginine biosynthesis complementation enables larval settlement in a marine sponge

**DOI:** 10.1101/2020.08.06.240770

**Authors:** Hao Song, Olivia H Hewitt, Sandie M Degnan

## Abstract

Larval settlement and metamorphosis are regulated by nitric oxide (NO) signalling in a wide diversity of marine invertebrates (1-10). It is surprising, then, that in most invertebrates, the substrate for NO synthesis – arginine – cannot be biosynthesized but instead must be exogenously sourced (11). In the sponge *Amphimedon queenslandica*, vertically-inherited proteobacterial symbionts in the larva are able to biosynthesize arginine (12,13). Here we test the hypothesis that symbionts might provide arginine to the sponge host so that nitric oxide synthase expressed in the larva can produce NO, which induces metamorphosis (8), and the byproduct citrulline (Fig. 1). First, we find support for an arginine-citrulline biosynthetic loop in this sponge larval holobiont using stable isotope tracing. In symbionts, incorporated ^13^C-citrulline decreases as ^13^C-arginine increases, consistent with the use of exogenous citrulline for arginine synthesis. In contrast, ^13^C-citrulline accumulates in larvae as ^13^C-arginine decreases, demonstrating the uptake of exogenous arginine and its conversion to NO and citrulline. Second, we show that while *Amphimedon* larvae can derive arginine directly from seawater, normal settlement and metamorphosis can occur in artificial sea water lacking arginine. Together, these results support holobiont complementation of the arginine-citrulline loop and NO biosynthesis in *Amphimedon* larvae, suggesting a critical role for bacterial symbionts in the development of this marine sponge. Given that NO regulates settlement and metamorphosis in diverse animal phyla (1-10) and arginine is procured externally in most animals (11), we propose that symbionts may play a equally critical regulatory role in this essential life cycle transition in other metazoans.

## Background, Results and Discussion

Larval settlement and metamorphosis are required for most marine invertebrates to complete their biphasic life cycle. These steps typically are initiated by the detection of environmental cues that indicate suitable habitat, and which in turn triggers signalling cascades that lead to behavioural and morphogenetic changes (3,14,15). Nitric oxide (NO) is a gaseous signalling molecule that plays a conserved role in regulating larval settlement and metamorphosis in a diverse range of marine invertebrates, including ascidians, gastropod and bivalve molluscs, polychaete annelids, bryozoans, cirripedian crustaceans and demosponges, as either an inhibitor (1-4,9,10) or an activator (5-8). In the marine sponge *Amphimedon queenslandica*, NO signalling induces larval settlement and metamorphosis (8) (Fig. 1A). However, arginine, which is a necessary precursor for NO synthesis, cannot be synthesized by *A. queenslandica*. Specifically, the genome of this sponge is missing genes encoding two necessary synthetases, namely argininosuccinate synthase, which converts citrulline into argininosuccinate, and argininosuccinate lyase, which catalyses arginine synthesis from argininosuccinate (16), although the gene encoded nitric oxide synthase (NOS) is present (Fig. 1B).

**Figure 1.**
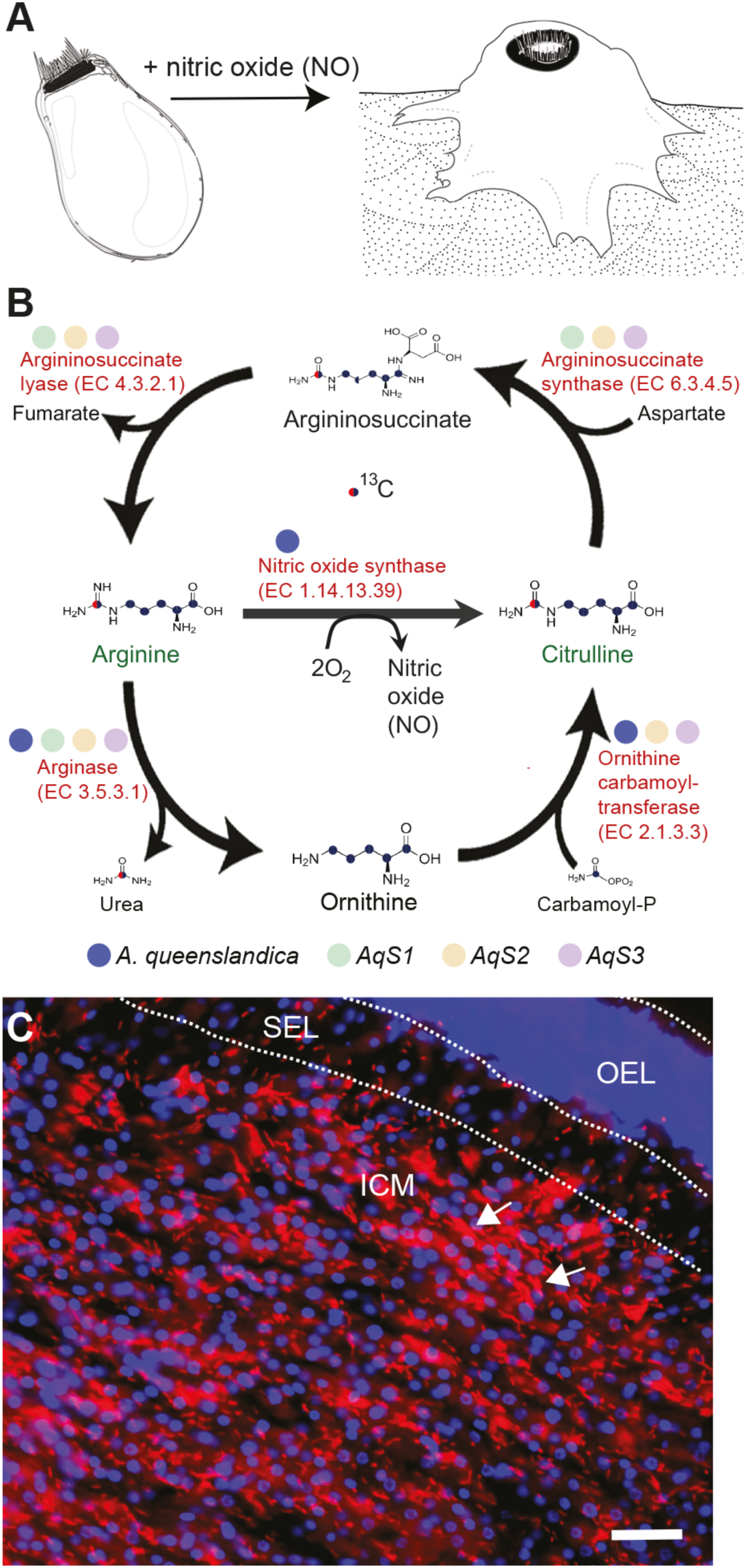
*Amphimedon queenslandica* cannot synthesize arginine, which is required for nitric oxide-induced settlement and metamorphosis. **A**. In *A. queenslandica*, nitric oxide (NO) is necessary for completion of larval settlement (Ueda et al.). **B**. Arginine, which is a necessary precursor for NO synthesis, cannot be synthesized the sponge. Its genome does not code for argininosuccinate synthase and argininosuccinate lyase, but both genes are present in its primary bacterial symbionts. Colored dots next to the enzymes comprising the arginine-citrulline synthesis loop correspond to presence in sponge and symbiont genomes. **C**. Fluorescence in situ hybridization (FISH) with an *AqS1*-specific 16S probe showing the distribution and abundance of this symbiont in the larval anterior. The outer epithelial layer (OEL), middle subepithelial layer (SEL) and inner cell mass (ICM) are shown. This symbiont is enriched in the inner cell mass (red; arrows pointing to examples); blue, DAPI-stained sponge cell nuclei. Scale bar 10 μm.

Although *A. queenslandica* adults can acquire arginine and other essential amino acids through their normal heterotrophic filter-feeding, sponge larvae are lecithotrophic (non-feeding) and therefore require another source of arginine to produce the NO necessary for settlement and metamorphosis. Potential sources of arginine in these non-feeding larvae are direct absorption from seawater and/or from symbiotic bacteria. *A. queenslandica* harbours a low complexity but stable bacterial community dominated by three proteobacterial symbionts – *AqS1, AqS2* and *AqS3 –* that are vertically transmitted with high fidelity from adults to embryos (12). All are present as extracellular symbionts in the inner cell mass of the larvae, with *AqS1* especially abundant (12) (Fig. 1C). Unlike *A. queenslandica*, the genomes of *AqS1, AqS2* and *AqS3* indicate they are capable of synthesizing arginine from citrulline (13) (Fig. 1B), and thus have the potential to supply their sponge larval host with the arginine needed for NOS to produce NO.

### *Amphimedon* larval settlement occurs in arginine-free sea water

Despite *A. queenslandica* lacking a complete arginine biosynthesis pathway, arginine is present in the free amino acid pool of multiple stages of the sponge life cycle, and is the most abundant free amino acid in the adult (Fig. S1). In swimming larvae, average arginine concentrations range from 0.053 nmol in larvae recently emerged from the adult (0-2 hours post emergence, hpe) to 0.043 nmol in larvae that are competent to settle (4-6 hpe; Fig. 2A). In comparison, arginine levels in natural seawater (collected from the native habitat of the sponge) and in artificial seawater (made from commercial sea salt), are 0.027 and 0.025 nmol mL^-1^, respectively (Fig. 2A). This means that approximately 1.75 ml of sea water could theoretically supply the total arginine present in a single larva.

**Figure 2.**
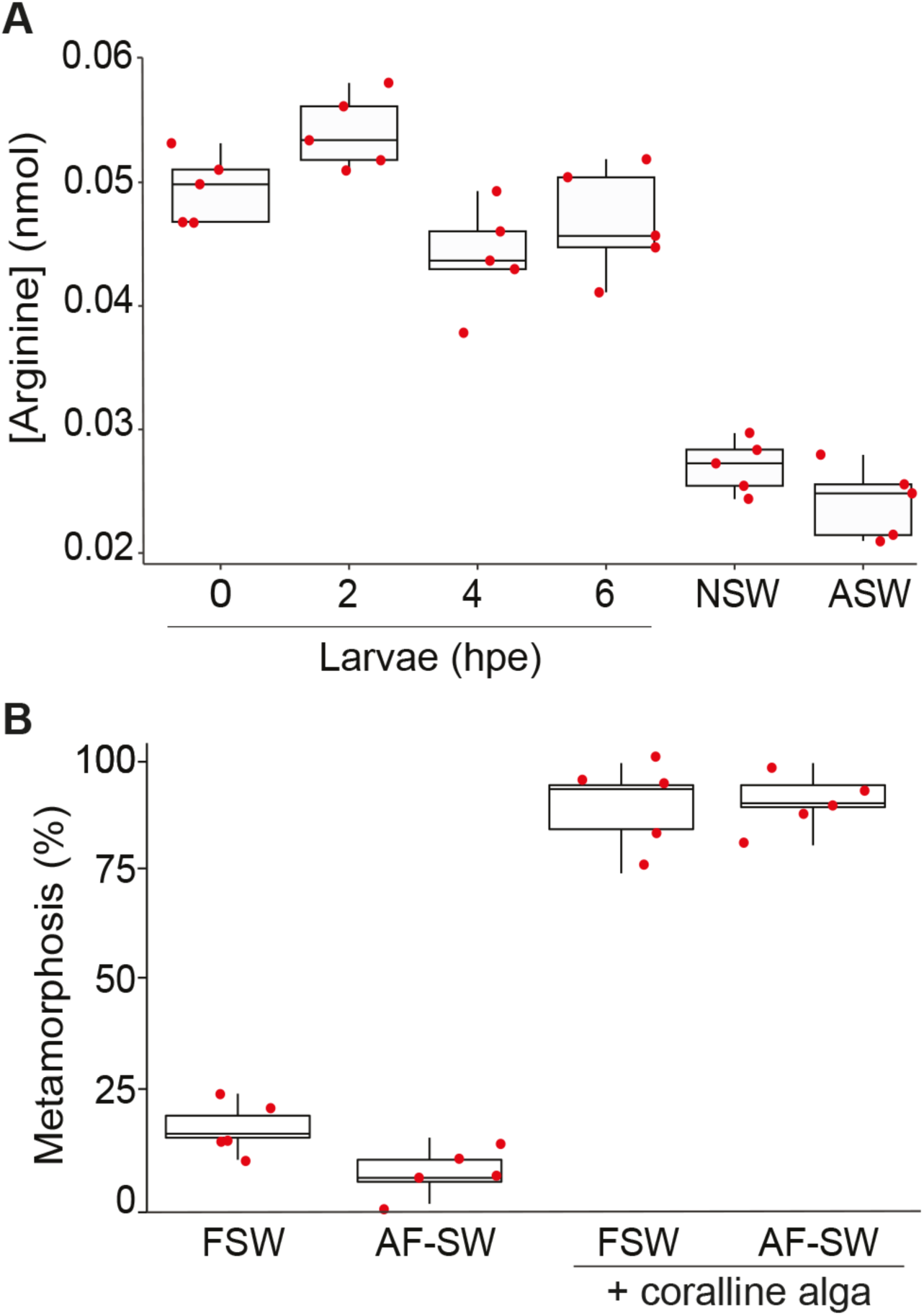
Exogenous arginine is not required for *Amphimedon queenslandica* larvae to settle and initiate metamorphosis. **A**. Arginine levels in individual larvae sampled at 0, 2, 4 and 6 hours post-emergence (hpe) from adult sponges in their native habitat on Heron Island Reef, and in 1 ml of natural sea water from Heron Island Reef (NSW) and of artificial seawater (ASW) made from commercial sea salts. Red dots, arginine levels in individual larvae or replicated water samples (n = 5). **B**. Percent of larvae that settled and initiated metamorphosis when reared in filtered sea water (FSW) or in arginine-free sea water (AF-SW), in the absence or presence of the inductive coralline alga *Amphiroa fragilissima* (15). Data are presented as the mean percentage of individuals that settled and initiated metamorphosis ± SEM. Red dots, 5 replicates (20 larvae per replicate).

We tested whether larvae rely on this uptake of exogenous arginine from sea water to undergo settlement and metamorphosis. To do so, we reared larvae in artificial seawater that contained no amino acids, and thus no arginine (arginine free seawater, AF-SW). We found that larvae reared in AF-SW were still able to successfully settle and metamorphose when induced by coralline algae, and could do so at a rate similar to those reared in natural seawater (Fig. 2B). Given that the induction of settlement and metamorphosis by NO signalling (8) can only occur in the presence of arginine, this result suggests that *A. queenslandica* larvae are accessing arginine from a source other than the surrounding sea water to drive NO production. We hypothesised that this source could be the bacterial symbionts.

### Evidence of holobiont complementation of NO biosynthesis in *Amphimedon* larvae

To test whether the symbiotic bacteria could be provisioning arginine for *A. queenslandica* larvae, we performed stable isotope-labelling with ^13^C-arginine and ^13^C-citrulline separately on sponge larvae and on bacterial symbionts. First, we incubated larvae in 10 μM ^13^C-labelled arginine in 0.22 μm-filtered seawater (FSW), and then measured the relative abundance of ^13^C-citrulline and ^13^C-arginine in these larvae at 1 and 5 h post incubation using high performance liquid chromatography (HPLC) and liquid-chromatography-mass spectrometry (LC-MS) analysis. Although the ratio of ^13^C-arginine/^12^C-arginine did not change over this period (p=0.130), we found a significant 5.7-fold increase in the ratios of both ^13^C-citrulline/^12^C-citrulline (p=0.005) and ^13^C-citrulline/^13^C-arginine (p=0.043) (Fig. 3). Since no ^13^C-citrulline was added in this experiment, the observed increase of ^13^C-citrulline must come from the labelled arginine; this demonstrates that the sponge larval holobiont can uptake arginine to produce NO and citrulline (Fig. 3). We observed a net arginine influx rate of ∽9.72 fmol larva^-1^ h^-1^ in the *A. queenslandica* larval holobiont, which is comparable to uptake rates of arginine and other free amino acids reported, for example, in *Crassostrea gigas* larvae (17).

**Figure 3.**
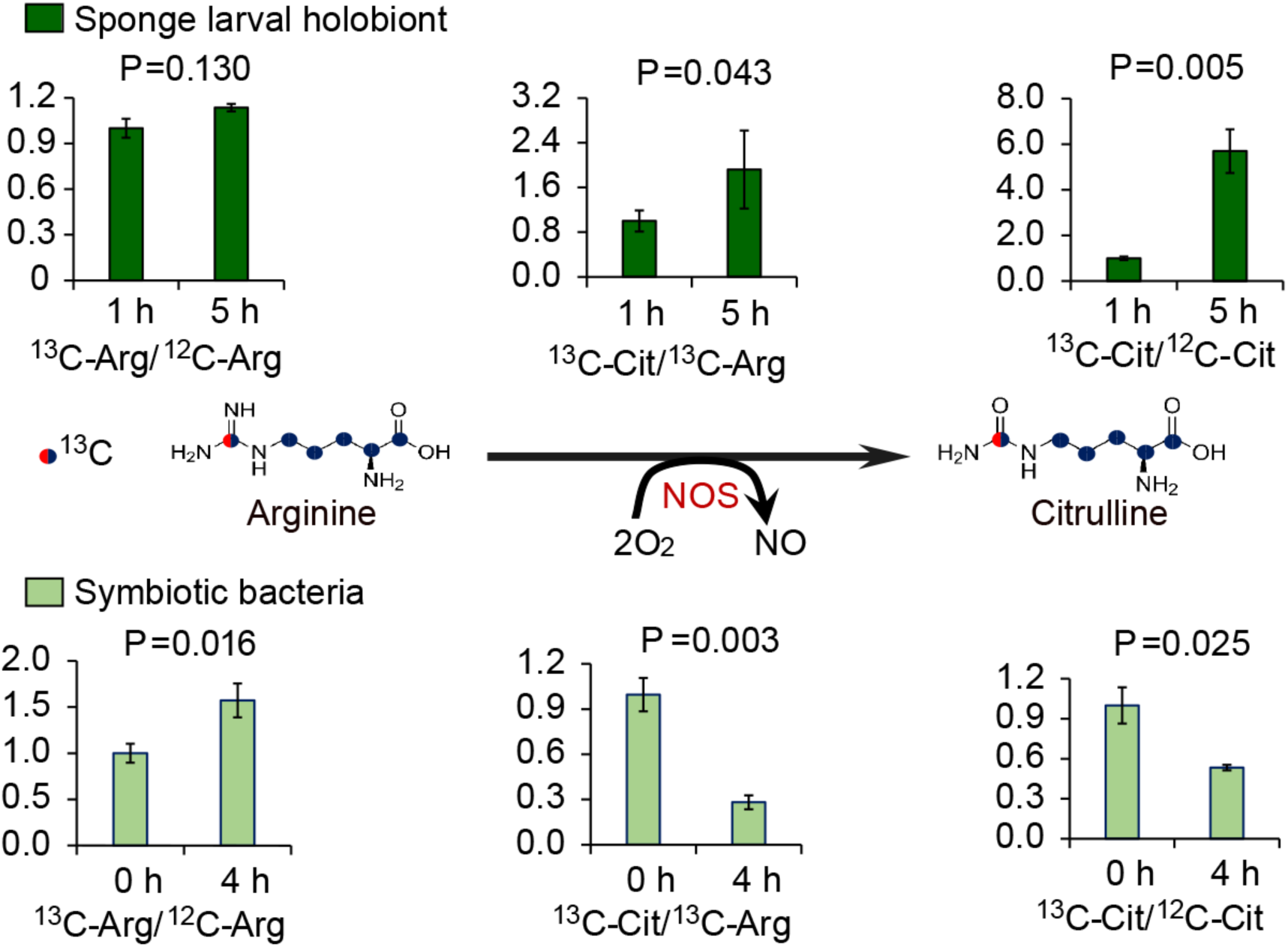
Evidence for arginine biosynthesis complementation between the sponge host and its bacterial symbionts. Relative changes in abundance of ^13^C in arginine and citrulline in *A. queenslandica* and its symbionts. *A. queenslandica* larvae were incubated with ^13^C–arginine (dark green, top row) for 1 and 5 h and the abundance of ^12^C- and ^13^C-citrulline, and ^12^C- and ^13^C-arginine were measured. The ordinate shows the relative change in abundance, where abundance at 1 h is 1.0. Symbiotic bacteria were incubated with ^13^C –citrulline (light green, bottom row) for 1 h and the abundance of ^12^C- and ^13^C-citrulline, and ^12^C- and ^13^C-arginine were measured at 1 and 4 h. The ordinate shows the relative change in abundance, where abundance at 0 h is 1.0.

Since citrulline can be synthesized by *A. queenslandica* (Fig. 1A), we next determined if the symbiotic bacteria can potentially use host-derived citrulline to biosynthesize arginine. To do this, we incubated an enriched symbiotic bacterial cell preparation with 10 μM ^13^C-labelled citrulline in FSW for 1 h. We measured the relative abundance of citrulline and arginine in the bacteria at the time of introduction of ^13^C-labelled citrulline (0 h) as well as 4 h later. We found that the relative ratio of ^13^C-citrulline/^12^C-citrulline decreased by approximately 47% (p=0.025), while that of ^13^C-arginine/^12^C-arginine increased approximately 53% (p=0.016, Fig. 3). Concurrently, the relative ratio of ^13^C-citrulline/^13^C-arginine decreased approximately 70% (p=0.003). Together, these results indicate that the symbiotic bacteria were uptaking the exogenous citrulline, and that a large proportion of this incorporated citrulline was rapidly converted into arginine.

## Conclusions

Here we provide experimental evidence of metabolic complementation between *A. queenslandica* and its primary bacterial symbionts that appears necessary for settlement and metamorphosis, and thus completion of this sponge’s life cycle. Specifically, these results are consistent with *A. queenslandica* larvae obtaining arginine, which is necessary for NO-induced settlement and metamorphosis (8), from its vertically-inherited symbiotic bacteria. Importantly, the ability of *A. queenslandica* larvae to metamorphose in arginine-free sea water is consistent with the symbionts, rather than seawater, being the primary source of arginine for the holobiont.

There appears to be at least two mechanisms by which symbiont-derived arginine could be acquired by *A. queenslandica* larval cells. First, the *A. queenslandica* genome encodes two cationic amino acid transporters (CATs) (*Aqu2*.*1*.*32956_001* and *Aqu2*.*1*.*32959_001*; see http://metazoa.ensembl.org/Amphimedon_queenslandica/Info/Index) that are expressed in larvae and throughout the sponge life cycle (Fig. S2, data acquired from *A. queenslandica* developmental dataset NCBI accession <u>GSE54364</u>). Arginine synthesized by the symbiotic bacteria could be transported into sponge cells by these CATs. Second, sponge cells in larvae could acquire symbiont-derived arginine by phagocytosis and digestion of their symbiotic bacteria. Phagocytosis is a common mechanism by which animal host cells acquire nutrients from their symbionts (18), and occurs in diverse marine endosymbioses such as the annelid *Olavius algarvensis* (19) and the deep-sea clam *Calyptogena magnifica* (20,21). Given that we have frequently observed phagocytosis of the dominant bacterial symbionts in *A. queenslandica* larvae and early postlarvae (12), we propose this as another potential pathway by which the sponge larva can acquire arginine from its symbiotic bacteria.

NO regulates settlement and metamorphosis in diverse marine invertebrates, including species of ascidians, gastropod and bivalve molluscs, polychaete annelids, bryozoans, cirripedian crustaceans and demosponges (1-10). NO can either repress or activate metamorphosis depending on the species, and these opposing activities are correlated neither with larval type (i.e. non-feeding lecithotrophs vs feeding planktotrophs) nor with phylogenetic position in the animal kingdom. Amongst these marine invertebrates, the arginine biosynthesis pathway is known only for the ascidian *Ciona intestinalis* (11) and *A. queenslandica* (this study), and in both cases appears to be incomplete. The same is true for the model invertebrates the fruit fly *Drosophila melanogaster*, the mosquito *Anopheles gambiae* and the nematode *Caenorhabditis elegans* (11). Given the apparent widespread requirement for an exogenous source of arginine for the biosynthesis of NO, and the role of NO in regulating settlement and metamorphosis, our study suggests that production of arginine by symbionts may be necessary for the completion of a diversity of marine invertebrate life cycles.

## Acknowledgements

The authors thank Manuel Plan, Gert Talbo and Terra Stark from the Australian Institute for Bioengineering & Nanotechnology at the University of Queensland for assistance in HPLC and LC-MS analysis, Davide Poli for guidance on bacterial cell enrichments, and Laura Rix for editorial assistance on an early draft of the manuscript. Heron Island Research Station provided logistical support for collection of animals. This research was funded by Australian Research Council grants DP110104601 and DP170102353 to S.M.D., and a Natural Science Foundation of Shandong Province travel grant ZR2019BD003, a China Postdoctoral Science Foundation grant 2019M652498 and China Scholarship Council visiting scholar grant to H.S.

## Author Contributions

S.M.D. and H.S. conceptualized this project and the methodological strategies. H.S. conducted all of the isotope and larval settlement experimental work and data analysis, with advice from S.M.D. O.H. conducted the FISH for bacterial symbionts. H.S. prepared the original draft of text and figures, and S.M.D. made significant contributions to revising the text and figures. All authors read and approved the final draft.

## Methods

### Larval collection

*Amphimedon queenslandica* sponge adults were collected from Heron Island Reef, southern Great Barrier Reef, Australia (23° 27’ S, 151° 56’ E) and maintained in flow-through ambient sea water aquaria at Heron Island Research Station. Adult sponges were induced to release larvae by mild heat treatment (1–2 °C above ambient temperature) for less than 2 h as previously described (22) and maintained in 500 ml native seawater at ambient temperature and light until use in experiments.

### Fluorescence in situ hybridization (FISH)

*A. queenslandica* larvae were fixed and stored in 70% ethanol, and embedded in paraffin wax for sectioning as described in 22 and 23. Embedded larvae were sectioned longitudinally to a thickness of 6 μm and mounted. Prior to hybridisation, larval sections were rehydrated in a graded series (70-20% ethanol, 1X PBS) and rinsed with hybridisation buffer [HB: 0.9 M NaCl, 20 mM Tris-HCl (pH 7.2), 0.01% SDS, 18% formamide (as described in 24). To visualise bacterial symbionts by FISH, we used the specifically designed oligonucleotide for *AqS1* (5′CCCCAGAGTTCCCGGCCGAA-3’, 12) that was 5’ end labelled with the indocarbocyanine fluorochrome, Cy3 (SigmaAldrich). This probe was dissolved in Ultrapure distilled water (Invitrogen) to 50 ng μL^−1^. 8 μl HB was mixed with 1 μl Cy3-labelled probe and applied to individual tissue sections, and incubated for 3 h at 46°C. Following hybridisation, samples were washed for 10 min at 48°C in prewarmed wash buffer [225 mM NaCl, 20 mM Tris-HCl (pH 7.2), 0.01% SDS (24)]. As a counter stain to enable visualisation of sponge cells, larval sections were counterstained with DAPI (4′,6diamidino-2-phenylindole, Sigma-Aldrich; 1:1000 μL PBS dilution) and incubated for 15 min in the dark at room temperature. Slides were rinsed with 1X PBS and fresh water, dried, then mounted in ProLong Gold antifade reagent (Life Technologies) and cured in darkness overnight before visual analysis. FISH and DAPI stained bacterial and sponge cells were visualised using confocal microscopy.

### Free amino acid extraction from sponge larvae, sponge adults and sea water

For free amino acid extractions, larvae were sampled at 0 h post-emergence from the maternal adult sponge (hpe), and then again at 2, 4 and 6 hpe. At each time point, 10 larvae were placed in 800 μl of chilled 80% ethanol and 1 μl of the internal standard norvaline (5 nmol μl^-1^ in water) (25) was added; this was replicated 5 times for each time point. Samples were then frozen at -80°C until further processing. For free amino acid extraction from adult sponges, ∽ 4 cm^3^ tissue was dissected by scalpel from 4 individuals under sterile conditions, and gently squeezed to remove excess seawater. Tissue samples were weighed, 800 μl of chilled 80% ethanol and 1 μl norvaline solution were added, and samples were frozen at -80°C.

Larval and adult samples were homogenized with zirconia beads (2 mm) at 4000 rpm for 30 sec using a Micro Smash MS-100 cell disrupter (Tomy Seiko Co., Ltd., Tokyo, Japan) and the homogenate centrifuged at 19,880 *g* for 10 min at 4°C. The supernatant was dried using a vacuum centrifugal evaporator and then re-dissolved by in 10 μl 500 μM 1,2-amino butyric acid, 500 μM sarcosine and 10 μl 20 % acetonitrile, and then stored at -80°C until high-pressure liquid chromatography (HPLC) analysis.

800 μl sea water samples were procured from: (i) the reef flat of Heron Island Reef at ∽10 cm below sea surface at mid-tide (transferred back to Heron Island Research Station on ice); and (ii) artificial seawater made fromTrophic Marin Pro Reef sea salt mixed with reverse osmosis filtered tap water. For both sea water samples, 1 μl norvaline solution was added, followed by evaporation and dissolving as described above, then stored at -80°C until HPLC analysis.

Free amino acids were quantified using a high-throughput HPLC method as described in 26.

### Settlement and metamorphosis assays

Larval metamorphosis assays were performed in sterile 6-well plates following 15. Newly-emerged larvae were incubated continuously in either 10?mL 0.22 μm filtered sea water (FSW) or 10 mL amino acid-free artificial seawater (AF-SW, recipe from http://cshprotocols.cshlp.org/content/2012/2/pdb.rec068270.full). Each treatment consisted of 10 wells with 20 larvae per well. At 6 hpe, by which time all larvae would be competent to settle (27), freshly prepared shards of the inductive coralline algae *Amphiroa fragilissima* were added to five wells in both the FSW and AF-SW to induce settlement. The remaining 5 wells per treatment served as controls, with no inductive coralline algae added. Plates were kept in the dark at 25°C and the number of larvae settled and initiating metamorphosis was scored after 24 h.

### Enrichment and collection of symbiotic bacteria

To isolate symbiotic bacteria, ∽ 5 g wet weight of freshly collected adult sponge tissue was dissected by sterile scalpel, cut into pieces, placed into individual 50 ml falcon tubes containing ice cold calcium and magnesium free artificial seawater (CMF-SW) and agitated on a shaker table for 5 min at 200 rpm. The supernatant was discarded and the sponge tissue rinsed 5 times in CMF-SW. Sponge cells and bacterial symbionts were dissociated by squeezing the tissue with a syringe through 25 μm mesh. The dissociated cells were centrifuged at 100 *g* for 15 min at 4°C to separate the sponge cells (pellet) from the symbiont bacteria (supernatant). The supernatant containing the bacterial cells were transferred to a new 50 mL tube and re-centrifuged at 200 *g* for 15 min at 4°C, and this step was repeated for a final centrifugation at 300 *g* for 15 min at 4°C. The final supernatant was filtered through a 5 μm mesh, and the filtrate was centrifuged for 20 min at 8800 *g* and 4°C to pellet the bacterial cells. The supernatant was then discarded and the bacterial cell pellet re-suspended in FSW (28). The final suspension was confirmed by microscopy to consist of ∽98% bacterial cells.

### Stable isotope tracer experiments

Larvae collected at 0 hpe were separated into four 1.5 mL tubes each containing 10 individuals and incubated with 1.2 mL FSW + 1.2 μl 10 mM ^13^C-arginine (Lot No. PR-21659, Catalog No. CLM-2070-0, Cambridge Isotope Laboratories, Inc.). From each tube, 30 μl of supernatant was removed every 2 h and processed for amino acids extraction as described above. The supernatants were analysed by LC-MS to determine the ^13^C/^12^C-arginine changes in the FSW.

Additional batches of 0 hpe larvae were incubated with 1.2 mL FSW + 1.2 μl 10 mM ^13^C-arginine as above (n=4 replicates, 10 larvae per replicate). Larvae were collected 1 and 5 h post incubation and washed three times in 1.2 mL FSW (centrifuged at 100 *g* for 3 min) and processed for extraction of amino acids as described above. The extraction solutions were moved into vials by Zip-Tip C18 for LC-MS analysis to determine the relative ratios of ^13^C/^12^C-arginine and ^13^C/^12^C-citrulline.

Four aliquots of 200 μl of the bacterial suspension obtained as described above were added to 2 μl 10 mM ^13^C-citrulline (Lot No. PR-16536, Catalog No. CLM-4899-0, Cambridge Isotope Laboratories, Inc., Tewksbury, MA, USA). After 1 h of incubation, the bacteria were washed with FSW three times, centrifuged for 10 min at 8800 *g*, and re-suspended in 1 mL FSW. Bacterial samples were collected 0 and 4 h post incubation (n=3 replicate samples for each time point) by centrifuging at 8800 *g* for 10 min, and were processed for amino acid extraction as described above. The extracted solutions were moved into vials using Zip-Tip C18 (Millipore, Bedford, MA, USA) for gas chromatography-mass spectrometry (GC-MS) analysis to determine the relative ratios of ^13^C/^12^C-arginine and ^13^C/^12^C-citrulline.

### LC-MS

LC-MS was carried out to quantify the proportion of arginine and citrulline either labelled or unlabelled with ^13^C isotope. For LC-MS, 6 μl aliquots were injected onto an Acquity UPLC CSH column (300 μm×100 mm, C-18, 1.7 μm, 130 Å, Waters Corporation, Milford, MA, USA). The flowrate was 3 μl min^-1^ and the analytes were eluted using a gradient of ACN in H2O. Initially, 3 min at 1% ACN, then 60% for 10 min and 95% for 12 min, before returning to the initial 1% ACN for 20 sec and re-equilibration for a total run-time of 20 min. The eluted analytes were mass analysed using an Exactive HF-X (Thermo Fischer Scientific, Walford, MA, USA) in targeted positive ion SIM mode with a resolution of 240,000. Five ions were targeted: 118.08626 (norvaline), 175.11896 (arginine), 176.19296 (^13^C-arginine), 176.12762 (citrulline), 177.11163 (^13^C-citrulline). Data were manually annotated using the Thermo Xcalibur Qual Browser software (Thermofisher).

### Statistics

Data were analysed by one-way analysis of variance (ANOVA) with treatment as a factor in SPSS 22.0. Significant differences among treatments were detected by Tukey’s honest significant difference post-hoc test. P values below 0.05 were considered significant.

