## Supplementary material for "Bacterial symbionts in animal development: arginine biosynthesis complementation enables larval settlement in a marine sponge": Fig. S1

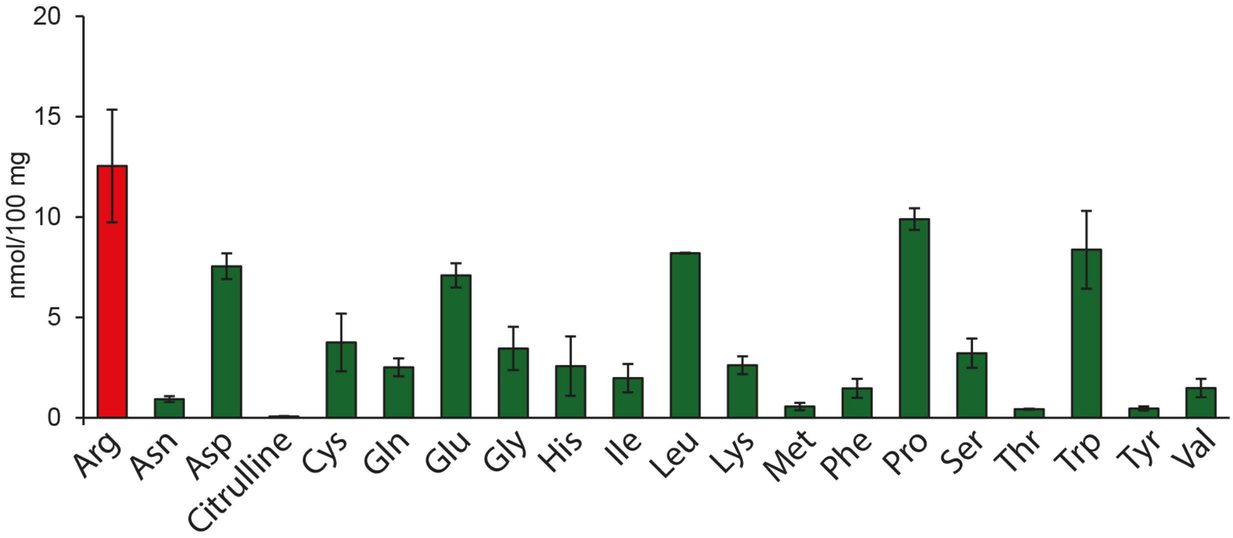


**Figure S1. Abundance of free amino acid levels in adult *A. queenslandica*.** Levels of each free amino acids was detected by HPLC, with arginine shown in red bar and others in green bar (n=4, mean and standard error are shown).
