## Supplementary material for "Bacterial symbionts in animal development: arginine biosynthesis complementation enables larval settlement in a marine sponge": Fig. S2

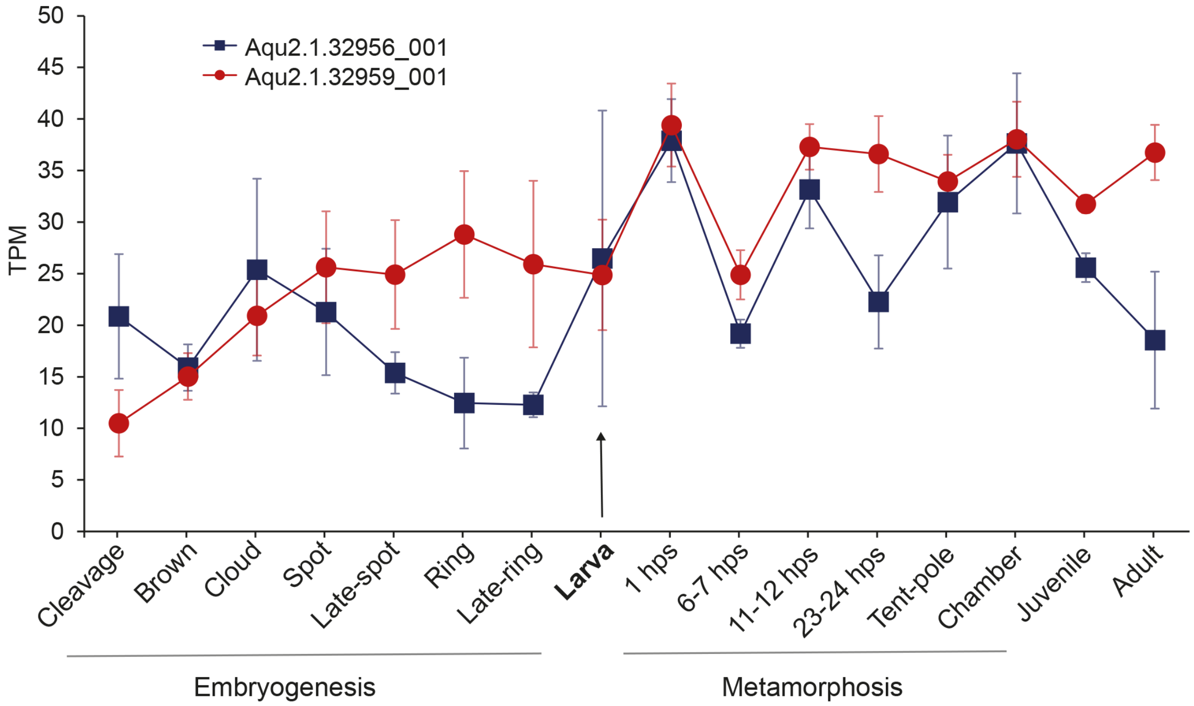


**Figure S2. Developmental expression profiles of two cationic amino acid transporters (CATs) (*Aqu2.1.32956_001* and *Aqu2.1.32959_001*).** CEL-Seq2 data are from 82 developmental samples from 15 stages of embryogenesis, larvae and metamorphosis, and adult *A. queenslandica* (NCBI accession GSE54364). Transcripts of Aqu2.1.32956_001 and Aqu2.1.32959_001 were observed in transcriptome datasets across multiple developmental stages with transcripts per million (TPM) varying from 2.2 to 95.3 and 2.3 to 51.6 respectively.
